# A direct SCN-to-DMH output pathway organizes circadian behavioral timing

**DOI:** 10.64898/2026.05.21.726959

**Authors:** Nicole E. Keene, Farina Pourmir, Ethan Hang, Mustafa Ozturget, Dylan A. McCreedy, Jeff R. Jones

## Abstract

Circadian rhythms in behavior depend on the suprachiasmatic nucleus (SCN), but how SCN timekeeping is transmitted to downstream circuits that organize daily behavioral rhythms remains poorly defined. The dorsomedial hypothalamus (DMH) has been implicated in circadian behavioral output, but lesion studies cannot determine whether the DMH contributes through local molecular timekeeping or through intact neurons that relay SCN-derived timing signals. Here, we combined DMH neuronal ablation, local molecular clock disruption, retrograde and intersectional tracing, single-cell RNA sequencing, and intersectional optogenetics to define and test a direct SCN-to-DMH output pathway. DMH neuronal ablation disrupted locomotor activity rhythms, whereas DMH *Cry1/2* disruption did not, indicating that intact DMH neurons, but not local molecular timekeeping, are required for locomotor rhythmicity. Retrograde tracing identified a sparse population of DMH-projecting SCN neurons concentrated in the dorsal SCN. These neurons showed minimal overlap with canonical AVP- and VIP-expressing SCN populations and were most strongly represented in a *Prokr2*/*Vipr2*-expressing transcriptional cluster. Repeated activation of DMH-projecting SCN neurons entrained locomotor rhythms, produced persistent phase shifts after stimulation ended, and compressed the temporal distribution of activity during entrainment. Together, these findings identify sparse, molecularly distinct DMH-projecting SCN neurons capable of organizing circadian locomotor timing.

## INTRODUCTION

Circadian rhythms organize daily patterns of physiology and behavior [1]. In mammals, these rhythms depend on the suprachiasmatic nucleus (SCN), the central circadian pacemaker. The mechanisms that generate and synchronize timekeeping within the SCN are well established, including molecular clock gene oscillations, neuronal synchrony across the SCN network, and entrainment by environmental light [2]. However, understanding how the SCN keeps time does not explain how that timing signal is routed to downstream neural circuits that organize behavior [3]. Identifying the pathways that carry SCN timing information beyond the SCN is therefore essential for linking central clock function to daily behavioral rhythms.

The dorsomedial hypothalamus (DMH) is a major hypothalamic output region involved in circadian regulation [4]. The DMH contributes to locomotor activity, arousal, feeding, corticosterone release, and sleep-wake regulation, all of which vary across the daily cycle [5–12]. Because locomotor activity rhythms are widely used as a behavioral assay of circadian organization, prior lesion studies have used this readout to show that disrupting the DMH severely impairs rhythmic behavioral output. These findings identify the DMH as an important node in the downstream control of circadian behavior, but they do not explain why intact DMH neurons are required for normal locomotor rhythmicity.

The same DMH lesion phenotype supports more than one mechanistic interpretation. Because SCN lesions also abolish locomotor rhythmicity, DMH-dependent locomotor activity rhythms cannot be interpreted as independent of SCN timing. Instead, the key distinction is whether DMH neurons require their own molecular clock to convert SCN-derived timing cues into rhythmic locomotor output [13]. Lesion studies cannot determine whether rhythmic output fails because local molecular timekeeping is disrupted or because DMH neurons are no longer available to receive and route SCN-derived signals [14,15]. Separating neuronal integrity from local clock function is therefore essential for defining how the DMH contributes to locomotor rhythmicity.

Timing information from the SCN could reach the DMH directly or indirectly [16–23]. SCN output to the DMH is commonly described as involving an indirect pathway through the subparaventricular zone (SPZ), consistent with the SPZ’s established role as an intermediate hypothalamic output region. However, this framework does not resolve whether the SCN also provides direct input to the DMH through a defined projection population. Anatomical studies have suggested direct SCN-to-DMH projections, but the location, molecular identity, and functional relevance of these neurons remain poorly defined.

The molecular identity of direct DMH-projecting SCN neurons cannot be inferred from anatomy alone. The SCN contains multiple neuropeptidergic populations, including arginine vasopressin (AVP)- and vasoactive intestinal peptide (VIP)-expressing neurons, which are enriched in dorsal and ventral SCN regions, respectively [24–27]. If a direct DMH-projecting SCN population were concentrated in the dorsal SCN, AVP neurons would be an obvious candidate source. However, projection-defined populations need not follow canonical neuropeptide boundaries [28,29]. Determining whether DMH-projecting SCN neurons overlap with AVP or VIP populations, or instead have a distinct transcriptional profile, is therefore necessary to define this output pathway.

Anatomical and molecular definitions of DMH-projecting SCN neurons do not by themselves establish whether this population can organize circadian behavioral output. The critical distinction is whether activating these neurons can reset the phase of the locomotor activity rhythm rather than produce only acute changes in activity [30]. If these neurons convey SCN output signals relevant to locomotor timing, repeated activation should impose a stable phase relationship between the stimulation interval and the behavioral rhythm and produce a persistent phase shift after stimulation ends. This distinction separates circadian entrainment from transient stimulation-evoked changes in activity [31].

Here, we tested whether the DMH requires local molecular timekeeping to support locomotor rhythmicity and whether direct SCN input to the DMH defines a functional circadian output pathway. We combined region-specific ablation and clock disruption, retrograde and intersectional tracing, single-cell RNA sequencing, and intersectional optogenetics to identify and functionally test a direct SCN-to-DMH output population. Together, these experiments identify DMH-projecting SCN neurons as a sparse, molecularly distinct subpopulation capable of organizing circadian locomotor timing.

## RESULTS

### DMH neuronal ablation impairs locomotor rhythmicity

To establish the behavioral consequences of neuronal loss in the SCN and DMH, we first genetically ablated neurons in each region by bilaterally co-injecting AAV-Syn-Cre with Cre-dependent Caspase-3 (Casp3) or diphtheria toxin A subunit (DTA) [32,33]. We then classified animals as lesion hits or misses based on post hoc histological verification of target-region cell loss (**Fig. 1a-c**). As expected, SCN-targeted hit animals lost free-running locomotor activity rhythms, whereas SCN-targeted miss animals remained rhythmic (**Fig. 1b**). Consistent with prior work implicating the DMH in the circadian control of locomotor activity, DMH-targeted hit animals showed severely dampened free-running rhythms, whereas DMH-targeted miss animals remained rhythmic (**Fig. 1c**). Locomotor activity rhythm amplitude, measured by Cosinor analysis, was significantly reduced in lesion hit animals relative to miss animals for both SCN- and DMH-targeted injections (two-way ANOVA with Fisher’s LSD post hoc test, p < 0.0001; **Fig. 1d**). Together, these results show that DMH neuronal loss markedly reduces locomotor activity rhythm amplitude.

**Figure 1.**
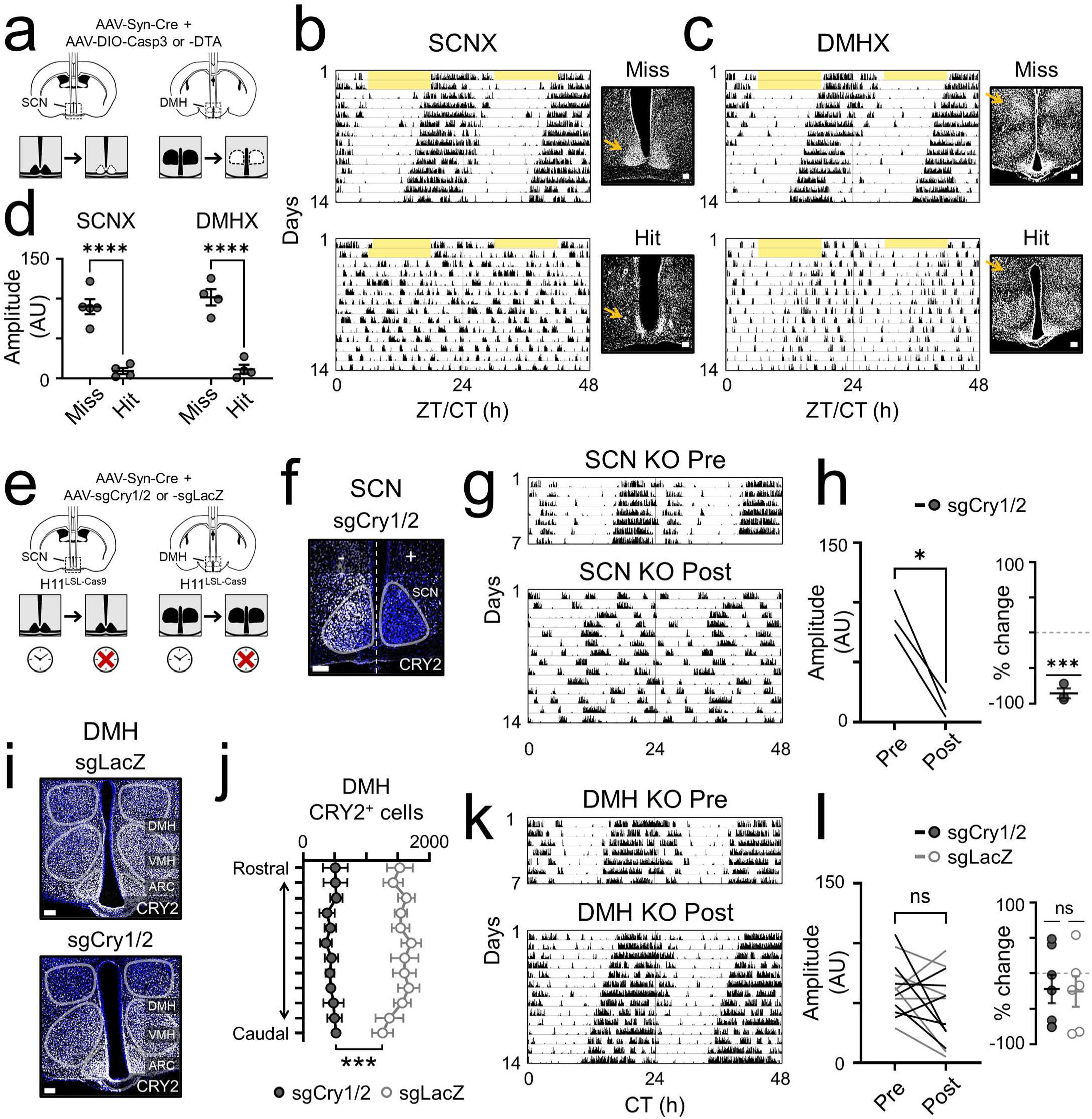
DMH ablation, but not clock disruption, impairs locomotor rhythmicity. **a)** Schematic of the genetic ablation strategy. AAV-Syn-Cre and AAV-DIO-Casp3 or AAV-DIO-DTA were co-injected bilaterally into either the suprachiasmatic nucleus (SCN) or dorsomedial hypothalamus (DMH) to genetically ablate neurons within the targeted region. **b,c)** Representative SCN- and DMH-targeted lesion hit/miss examples. In each panel, the top row shows an actogram and corresponding DAPI image from a lesion miss animal, and bottom row shows an actogram and corresponding DAPI image from a lesion hit animal. Yellow shading indicates lights on, black ticks indicate locomotor activity, and actograms depict 2 d in a 12 h:12 h light:dark cycle (LD) followed by 12 d in constant darkness (DD). **b)** SCNX, SCN ablation. SCNX miss animals preserved free-running locomotor activity rhythms, whereas SCNX hit animals lost locomotor rhythmicity. **c)** DMHX, DMH ablation. DMHX miss animals preserved free-running locomotor activity rhythms, whereas DMHX hit animals showed severely dampened rhythms. Yellow arrows indicate intact or lesioned SCN or DMH regions. **d)** Cosinor amplitude of locomotor rhythms in SCNX and DMHX hit and miss animals. Lesion hit animals showed significantly reduced rhythm amplitude relative to miss animals for both targets (two-way ANOVA with Fisher’s LSD post hoc test, p < 0.0001; n = 4-5 animals per group). **e)** Schematic of the genetic clock-disruption strategy. AAV-Syn-Cre and AAV-sgCry1/2 or control AAV-sgLacZ were injected into either the SCN or DMH to disrupt the local molecular clock without ablating the targeted neurons. **f-h)** SCN-targeted clock disruption. **f)** Representative unilateral SCN-targeted sgCry1/2 injection showing preserved CRY2 immunoreactivity in the non-targeted side of the SCN (sgCry1/2 -) and loss of CRY2 immunoreactivity in the targeted side (sgCry1/2+). **g)** Representative double-plotted actograms from the same animal before (7 d, DD) and after (14 d, DD) bilateral sgCry1/2 injection into the SCN, showing loss of locomotor rhythmicity after injection. **h)** Quantification of locomotor rhythm amplitude before and after SCN clock disruption. *Left*, Cosinor amplitude was significantly reduced after sgCry1/2 injection (paired t-test, p = 0.024). *Right*, percent change in Cosinor amplitude relative to pre-injection baseline was significantly reduced (one-sample t-test, p = 0.001; n = 3 mice). **i-l)** DMH-targeted clock disruption. **i)** Representative bilateral DMH-targeted sgLacZ (*top*) or sgCry1/2 (*bottom*) injection. CRY2 immunoreactivity is preserved throughout sgLacZ-targeted tissue and selectively lost from the sgCry1/2-targeted DMH, while remaining present in adjacent ventromedial hypothalamus (VMH) and arcuate nucleus (ARC). **j)** Quantification of CRY2-positive cells across the rostral-caudal extent of the DMH in sgLacZ- and sgCry1/2-targeted animals (40 µm coronal sections; n = 4-5 mice per group). CRY2-positive cells were significantly reduced in sgCry1/2 animals across the DMH (two-way repeated-measures ANOVA, significant main effect of genotype, p < 0.0001; no significant effect of slice or genotype × slice interaction). **k)** Representative double-plotted actograms from the same animal before (7 d, DD) and after (14 d, DD) bilateral sgCry1/2 injection into the DMH, showing preserved locomotor rhythmicity after injection. **l)** Quantification of locomotor rhythm amplitude before and after DMH clock disruption. Cosinor amplitude was not significantly altered after DMH-targeted sgCry1/2 or sgLacZ injection (two-way repeated-measures ANOVA with Fisher’s LSD post hoc test, p > 0.05), and percent change in Cosinor amplitude did not differ between sgCry1/2- and sgLacZ-injected animals (unpaired t-test, p > 0.05; n = 6-7 mice per group). In all images, scale bars = 100 µm.*, p < 0.05; ***, p < 0.001; ****, p < 0.0001; ns, not significant.

### DMH clock disruption does not impair locomotor rhythmicity

To determine whether the DMH ablation phenotype depended on the local molecular clock rather than neuronal loss itself, we bilaterally co-injected AAV-Syn-Cre with AAV-sgCry1/2 into mice expressing Cre-dependent Cas9 (H11^LSL-Cas9^; **Fig. 1e**) [34,35]. We first tested this strategy in the SCN, where disruption of *Cry1* and *Cry2* is expected to eliminate behavioral circadian rhythms. Consistent with prior work, SCN-targeted sgCry1/2 eliminated CRY2 immunoreactivity in targeted SCN neurons and abolished free-running locomotor activity rhythms following bilateral injection (**Fig. 1f,g**). After SCN clock disruption, locomotor activity rhythm amplitude was significantly reduced both as absolute Cosinor amplitude and as percent change from pre-injection baseline (paired t-test, p = 0.024; one-sample t-test, p = 0.001; **Fig. 1h**).

We then applied the same *Cry1*/2-disruption strategy to the DMH. DMH-targeted sgCry1/2 selectively eliminated CRY2 immunoreactivity across the rostrocaudal extent of the DMH while preserving CRY2 labeling in the adjacent ventromedial hypothalamus and arcuate nucleus (VMH and ARC; two-way ANOVA, significant main effect of genotype, p < 0.0001; no significant effect of slice or genotype × slice interaction; **Fig. 1i,j**). Even with robust CRY2 loss in the DMH, animals maintained free-running locomotor activity rhythms (**Fig. 1k**). In DMH-targeted animals, sgCry1/2 did not significantly alter locomotor activity rhythm amplitude relative to control sgLacZ, either as absolute Cosinor amplitude or as percent change from pre-injection baseline (two-way repeated measures ANOVA with Fisher’s LSD post hoc test, p > 0.05; unpaired t-test for percent change, p > 0.05; **Fig. 1l**). Together, these results show that DMH neuronal loss and DMH molecular clock disruption have dissociable effects: DMH ablation impairs locomotor activity rhythms, but local DMH clock disruption does not.

### The SCN contains a direct DMH-projecting population concentrated in the dorsal SCN

The dissociation between DMH neuronal ablation and DMH clock disruption suggested that the DMH’s contribution to locomotor rhythmicity depends on circuit integrity rather than local molecular timekeeping. Because SCN clock disruption abolished free-running locomotor activity rhythms, we next asked whether SCN neurons provide direct input to the DMH. To test this, we injected AAVrg-Syn-EGFP unilaterally into the DMH to label DMH-projecting neurons (**Fig. 2a**) [36]. Injection-site heatmaps showed broad DMH coverage across animals, with limited spread into the adjacent VMH and ARC (**Fig. 2b**).

**Figure 2.**
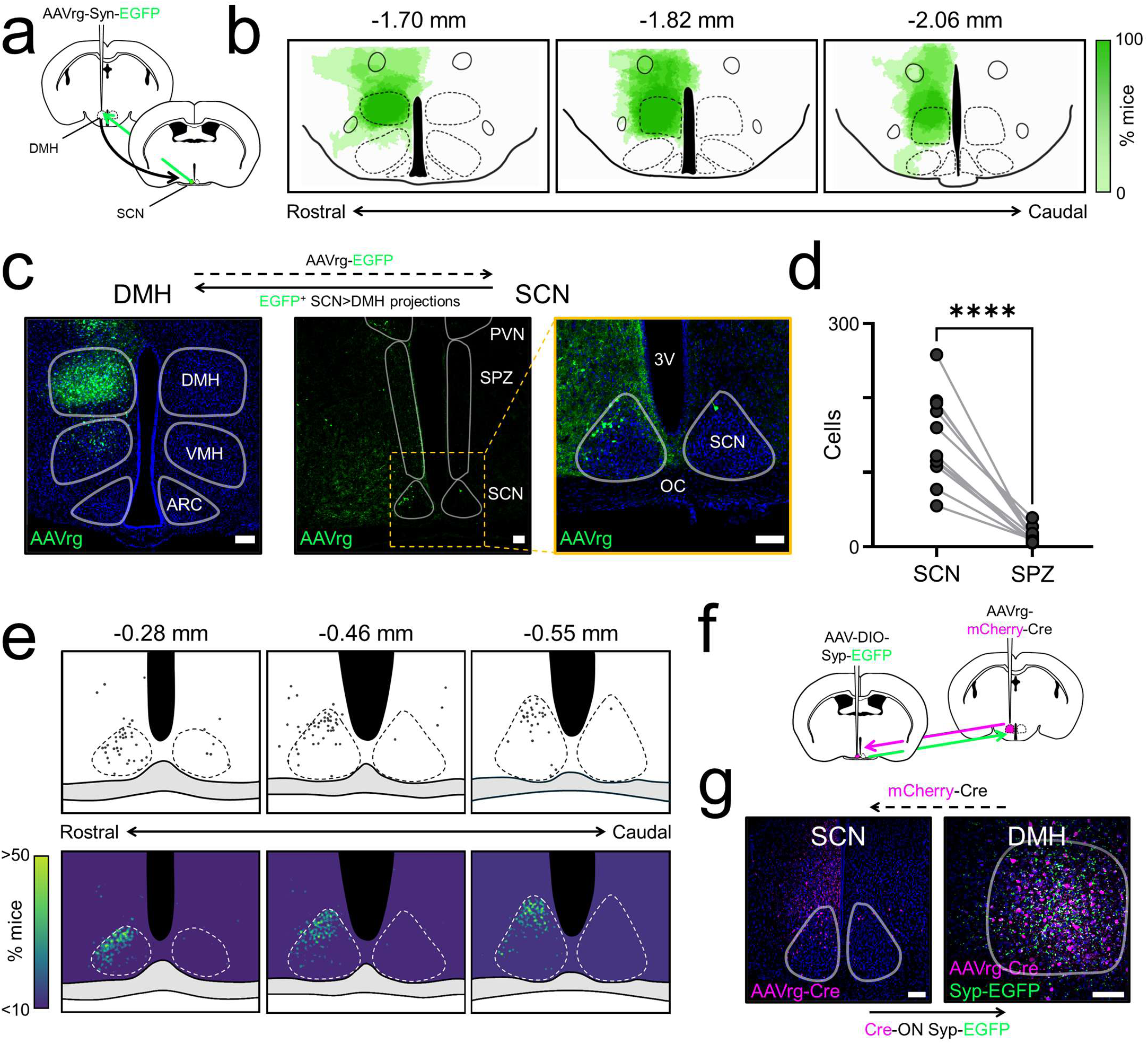
Direct SCN to DMH projection neurons are concentrated in the dorsal SCN. **a)** Schematic of the retrograde tracing strategy. AAVrg-Syn-EGFP was injected unilaterally into the dorsomedial hypothalamus (DMH) to DMH-projecting neurons, including neurons in the suprachiasmatic nucleus (SCN). **b)** Rostral-to-caudal atlas sections showing the extent of AAVrg-Syn-EGFP injection spread in the DMH across animals (n = 8). Green heatmap indicates the percentage of mice with injection coverage at each location, from low (light green) to high (dark green). White outlines indicate the DMH, ventromedial hypothalamus (VMH), and arcuate nucleus (ARC). **c)** Representative images of retrograde labeling after unilateral DMH-targeted AAVrg-Syn-EGFP injection. *Left*, injection site localized to the DMH. *Middle*, low-magnification image showing EGFP-labeled neurons concentrated in the SCN, with little labeling in adjacent regions. SPZ, subparaventricular zone; PVN, paraventricular nucleus. *Right*, higher-magnification image of the boxed SCN region showing EGFP-labeled SCN neurons. 3V, third ventricle; OC, optic chiasm. **d)** Quantification of retrogradely labeled SCN^DMH^ neurons in the SCN and adjacent SPZ. Significantly more EGFP-positive neurons were observed in the SCN than in the SPZ (paired t test, p < 0.0001; n = 8 mice). **e)** Distribution of SCN neurons projecting to the DMH. *Top*, representative rostral-to-caudal atlas sections from a single mouse showing the center of mass of each EGFP-positive cell within and around the SCN. *Bottom*, group heatmap across all animals (n = 8) showing the spatial distribution of EGFP-labeled DMH-projecting SCN neurons (SCN^DMH^), which were concentrated primarily in the dorsal SCN. Heatmap color indicates the percentage of mice with a labeled cell at a given location, from low (blue) to high (yellow). White outlines indicate the SCN. **f)** Schematic of the intersectional tracing strategy used to label synaptic terminals of SCN^DMH^ neurons. AAVrg-mCherry-Cre was injected into the DMH, and AAV-DIO-Syp-EGFP was injected into the SCN, enabling Cre-dependent synaptophysin-EGFP labeling of SCN neurons that project to the DMH. **g)** Representative images following the intersectional tracing strategy (n = 2 mice). *Left*, SCN showing retrogradely labeled mCherry-Cre-positive SCN^DMH^ neurons. *Right*, DMH showing mCherry-Cre labeling in the targeted DMH region together with synaptophysin-EGFP-labeled putative terminals from SCN^DMH^ neurons. In all images, scale bars = 100 µm. ****, p < 0.0001.

DMH-targeted AAVrg-Syn-EGFP labeled neurons concentrated in the SCN, with little labeling in adjacent regions such as the subparaventricular zone or paraventricular nucleus of the hypothalamus (SPZ or PVN; **Fig. 2c**). Because some injections extended dorsal to the DMH, we examined cases in which AAVrg-Syn-EGFP was centered above the DMH. These injections labeled cells dorsal to the SCN but did not label SCN neurons, indicating that SCN labeling in DMH-targeted animals was unlikely to result from viral uptake at sites dorsal to the DMH (**Supplementary Fig. 1**). Across animals, significantly more EGFP-positive neurons were located in the SCN than in the adjacent SPZ (paired t-test, p < 0.0001; **Fig. 2d**). We refer to these retrogradely labeled DMH-projecting SCN neurons as SCN^DMH^ neurons. Spatial mapping across rostrocaudal SCN sections showed that SCN^DMH^ neurons were concentrated primarily in the dorsal SCN (**Fig. 2e**). A full rostrocaudal image series showed that most labeled cells were within the SCN, with a small number of extra-SCN cells in rostral sections consistent with medial preoptic area or median preoptic nucleus populations that also project to the DMH (**Supplementary Fig. 2**). Together, these results identify a direct population of DMH-projecting SCN neurons concentrated in the dorsal SCN.

To test whether SCN^DMH^ neurons give rise to putative terminals in the DMH, we used an intersectional tracing strategy in which AAVrg-mCherry-Cre was injected into the DMH and Cre-dependent synaptophysin-EGFP (AAV-DIO-Syp-EGFP) was injected into the SCN (**Fig. 2f**) [37]. This approach labeled retrogradely identified SCN^DMH^ neurons in the SCN and synaptophysin-EGFP-positive putative terminals in the DMH (**Fig. 2g**). Thus, the retrogradely-defined SCN^DMH^ population provides an anatomical substrate for direct SCN input to the DMH.

### SCN^DMH^ neurons show minimal overlap with AVP and VIP populations

Because SCN^DMH^ neurons were concentrated in the dorsal SCN, where arginine vasopressin (AVP)-expressing neurons are enriched, we next asked whether this projection arose from AVP neurons or from another canonical SCN neuropeptide population, vasoactive intestinal peptide (VIP)-expressing neurons. To test this, we immunolabeled SCN sections containing retrogradely labeled SCN^DMH^ neurons for AVP and VIP (**Fig. 3a,b**). SCN^DMH^ neurons were sparse relative to both AVP- and VIP-positive neurons across the rostrocaudal extent of the SCN (**Fig. 3c**). Higher-magnification imaging showed that only a small subset of SCN^DMH^ neurons colocalized with AVP. Many SCN^DMH^ neurons were immediately adjacent to AVP-positive neurons but lacked detectable AVP coexpression (**Fig. 3d**). Population-level overlap analysis supported the same conclusion, with SCN^DMH^ neurons showing no detectable overlap with VIP and only limited overlap with AVP (**Fig. 3e-g**). Together, these results show that SCN^DMH^ neurons occupy the dorsal SCN near AVP-positive neurons but overlap only minimally with AVP- or VIP-expressing populations.

**Figure 3.**
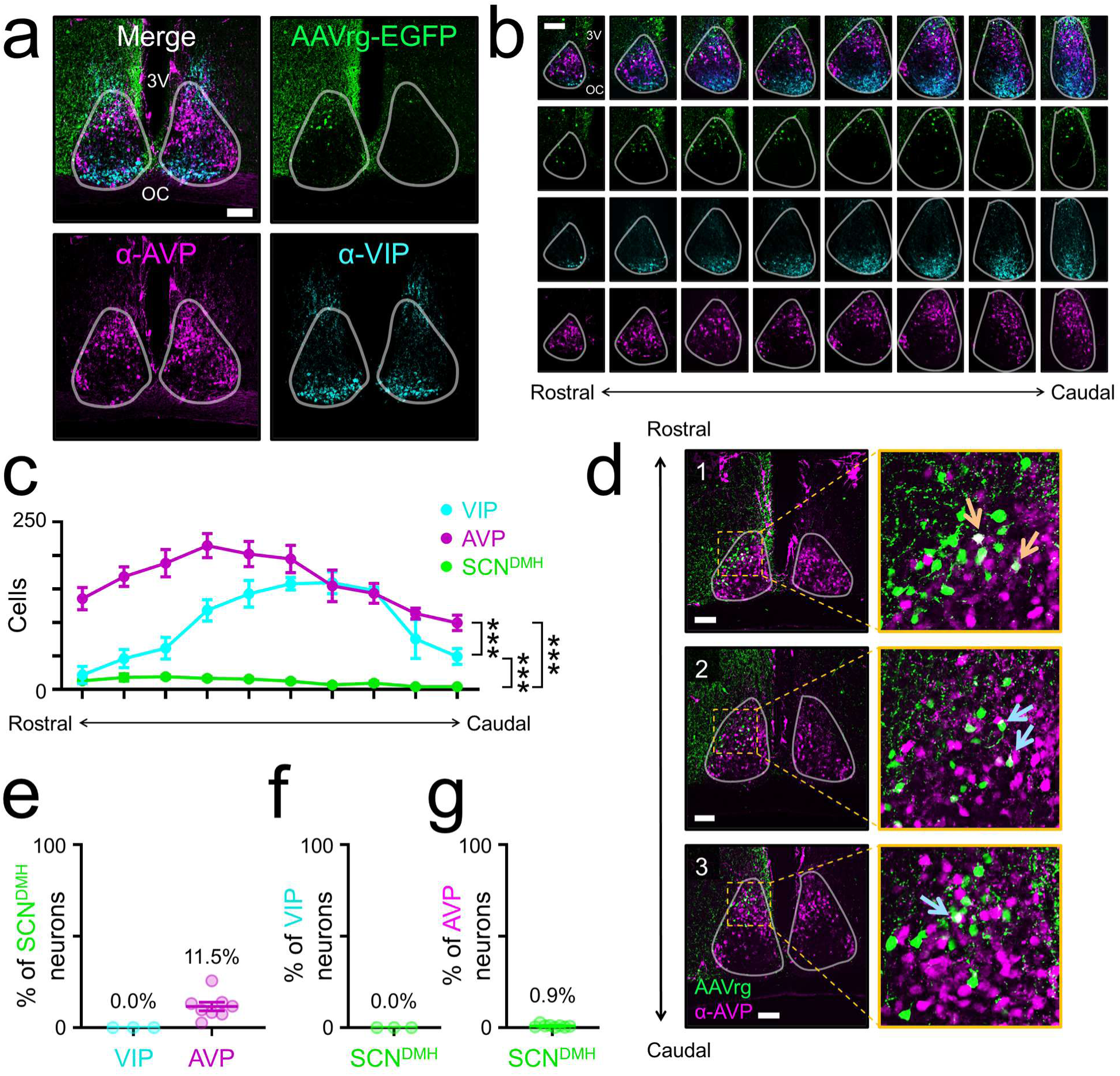
SCN^DMH^ neurons are sparse and show minimal overlap with AVP and VIP populations. **a)** Representative coronal SCN section showing retrogradely labeled SCN^DMH^ neurons (AAVrg-EGFP, green), AVP immunoreactivity (magenta), and VIP immunoreactivity (cyan). White outline indicates the SCN. 3V, third ventricle; OC, optic chiasm. **b)** Representative rostral-to-caudal series of SCN sections showing the distribution of SCN^DMH^ neurons (green), AVP-positive neurons (magenta), and VIP-positive neurons (cyan) across the SCN. **c)** Quantification of SCN^DMH^, AVP-positive, and VIP-positive neurons. Cell counts are plotted by 40 µm coronal SCN sections from rostral to caudal (SCN^DMH^, n = 8 mice; AVP, n = 8 mice; VIP, n = 3 mice; two-way ANOVA with Tukey’s multiple comparisons test, p <0.0001). **d)** Representative images from three mice showing SCN^DMH^ neurons (green) and AVP immunoreactivity (magenta). *Left*, low-magnification views of the SCN. Yellow dashed boxes indicate the regions shown at higher magnification at right. *Right*, higher-magnification views showing examples of SCN^DMH^ neurons that colocalize with AVP (orange arrows) and SCN^DMH^ neurons adjacent to, but not colocalizing with, AVP neurons (blue arrows). White outlines indicate the SCN. **e-g)** Quantification of overlap between SCN^DMH^ neurons and AVP or VIP populations. **e)** Percentage of SCN^DMH^ neurons co-expressing VIP or AVP. **f)** Percentage of VIP neurons co-labeled with EGFP. **g)** Percentage of AVP neurons co-labeled with EGFP. In all images, scale bars = 100 µm. ***, p < 0.001.

### SCN^DMH^ neurons are enriched in a transcriptionally distinct SCN cluster

The limited overlap with AVP and VIP suggested that canonical neuropeptidergic markers were insufficient to define the SCN^DMH^ population. We therefore used single-cell RNA sequencing to examine the molecular profile of retrogradely labeled SCN^DMH^ neurons beyond these markers. We injected AAVrg-Syn-EGFP into the DMH, microdissected and dissociated the SCN, and performed scRNA-seq with targeted EGFP transcript detection to identify retrogradely labeled neurons within the broader SCN population (**Fig. 4a**). Putative SCN neurons were isolated by manual curation after identifying the neuronal cluster and excluding non-SCN neuronal clusters enriched for *Sst* or *Trh* (**Supplementary Fig. 3**) [38]. We first asked whether EGFP-positive SCN^DMH^ neurons aligned with canonical SCN marker expression, including neuropeptide markers and receptors commonly used to define SCN neuron populations (*Avp*, *Vip*, *Cck*, *Nms*, *Vipr2*, *Grp*, and *Prokr2*) [28,29]. EGFP-positive SCN^DMH^ neurons were distributed across UMAP space rather than confined to a single marker-defined domain (**Fig. 4b**). Compared with the overall putative SCN neuron population, SCN^DMH^ neurons were relatively depleted for *Vip*, *Nms*, and *Grp*, relatively enriched for *Cck*, *Vipr2*, and *Prokr2*, and showed a similar fraction of *Avp*-positive cells (**Fig. 4c**).

**Figure 4.**
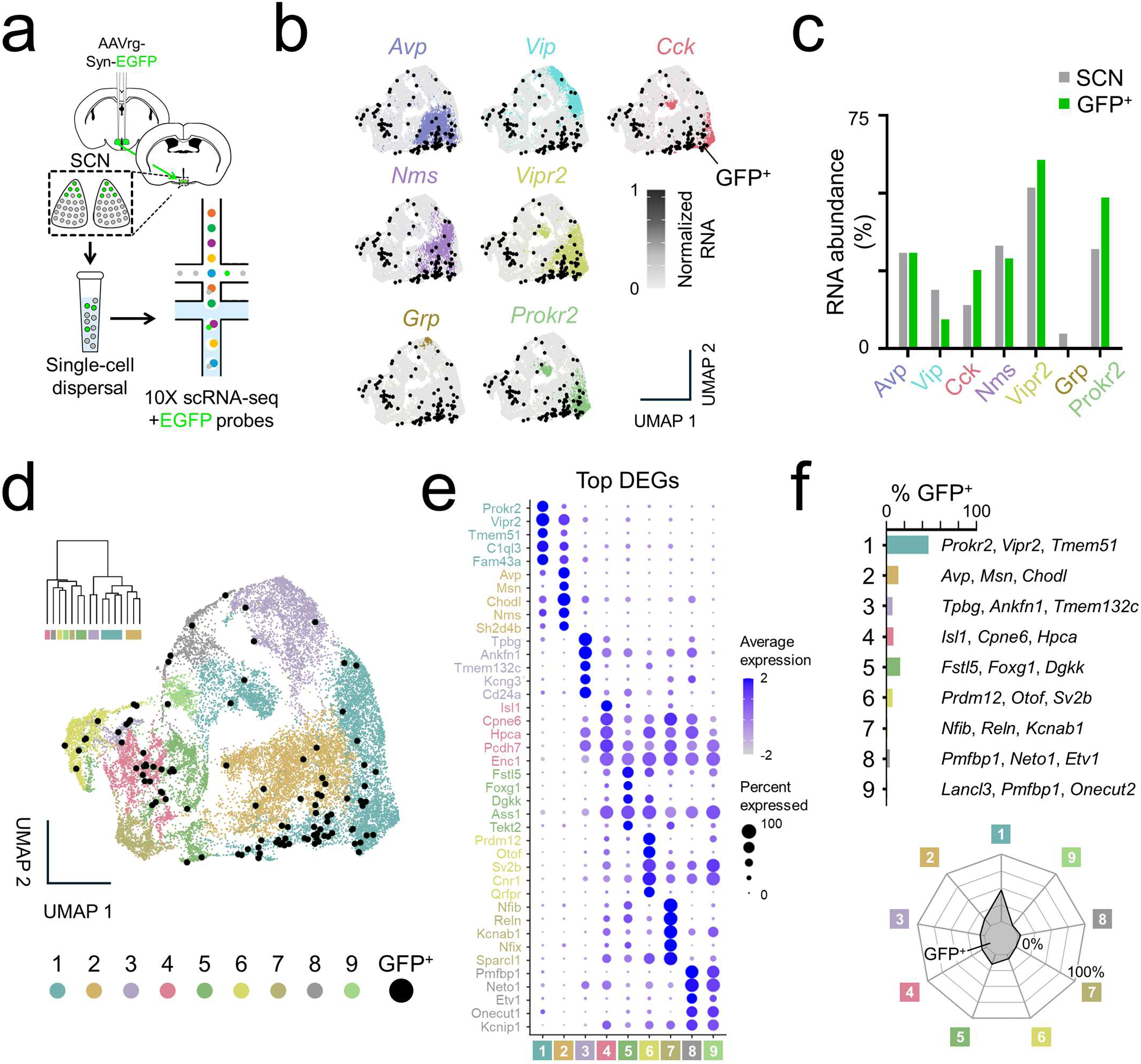
SCN^DMH^ neurons are a sparse population enriched in a *Prokr2*- and *Vipr2*-expressing cluster. **a)** Schematic of the scRNA-seq workflow. AAVrg-Syn-EGFP was injected bilaterally into the DMH to label DMH-projecting SCN neurons (SCN^DMH^). The SCN was then microdissected, dissociated into a single-cell suspension, and processed for 10x Genomics single-cell RNA sequencing with additional probes to detect EGFP transcripts, allowing identification of retrogradely labeled SCN^DMH^ neurons within the broader SCN cell population. **b)** UMAPs of putative SCN neurons showing normalized expression of canonical SCN marker genes (*Avp*, *Vip*, *Cck*, *Nms*, *Vipr2*, *Grp*, and *Prokr2*). Putative SCN neurons were identified by manual curation after initial isolation of the neuronal cluster and exclusion of non-SCN neuronal clusters enriched for *Sst* or *Trh* (see **Supplementary Fig. 3**). Black dots indicate EGFP-positive SCN^DMH^ neurons. **c)** Percentage of cells with detectable RNA expression (> 0) for canonical SCN marker genes in all putative SCN neurons (gray) and EGFP-positive SCN^DMH^ neurons (green). SCN^DMH^ neurons were relatively depleted for *Vip*, *Nms*, and *Grp*, relatively enriched for *Cck*, *Vipr2*, and *Prokr2*, and showed a fraction of *Avp*-positive cells similar to that of the overall SCN population. **d)** Unbiased clustering of putative SCN neurons. *Inset*, hierarchical clustering dendrogram generated by Ward linkage resolving nine clusters. *Main panel*, UMAP colored by cluster identity, with EGFP-positive SCN^DMH^ neurons shown in black. **e)** Dot plot showing the top five differentially expressed genes (DEGs) for each cluster. DEGs were identified using positive markers expressed in at least 25% of cells in the cluster with log_2_ fold change ≥ 0.25, then ranked within each cluster by adjusted p value; the top five genes per cluster are shown. Dot size indicates percent of cells expressing each gene, and color indicates average expression. **f)** Distribution of EGFP-positive SCN^DMH^ neurons across unbiased clusters. *Top*, percentage of all EGFP-positive neurons assigned to each cluster, with the top three DEGs for each cluster listed at right. *Bottom*, radar plot showing the same cluster distribution of EGFP-positive neurons. SCN^DMH^ neurons were concentrated in cluster 1, with smaller fractions in clusters 2 and 5 and the remainder distributed across the other clusters.

The broad distribution of SCN^DMH^ neurons across canonical marker domains led us to perform unbiased clustering of putative SCN neurons. Ward linkage hierarchical clustering resolved nine clusters, which were visualized on the UMAP and annotated using the top differentially expressed genes for each cluster (**Fig. 4d,e**) [39]. SCN^DMH^ neurons were sparse across the dataset and were distributed unevenly across clusters. The largest fraction of EGFP-positive SCN^DMH^ neurons was assigned to cluster 1, marked by *Prokr2*, *Vipr2*, and *Tmem51*, with smaller fractions in clusters 2 and 5 and the remaining cells sparsely distributed across other clusters (**Fig. 4f**). Together, these results show that SCN^DMH^ neurons are largely distinct from AVP and VIP populations and are enriched within a cluster marked by the canonical SCN genes *Prokr2* and *Vipr2*.

### Repeated stimulation of SCN^DMH^ neurons entrains locomotor activity rhythms

After defining SCN^DMH^ neurons anatomically and transcriptionally, we next tested whether repeated activation of these neurons is sufficient to entrain locomotor activity rhythms [40]. To selectively express the optogenetic activator channelrhodopsin (ChR2) in SCN^DMH^ neurons, we injected AAVrg-EGFP-Cre into the DMH and AAV-DIO-ChR2-mCherry into the SCN, then implanted an optical fiber above the SCN (**Fig. 5a,b**). Animals were maintained in constant darkness and received daily optogenetic stimulation for more than 2 weeks (470 nm, 10 Hz, 1 h/day). Representative SCN sections showed greater c-FOS immunoreactivity after stimulation in SCN^DMH^-ChR2 animals than in SCN EGFP controls, consistent with activation of ChR2-expressing SCN^DMH^ neurons by the stimulation protocol (**Fig. 5c**).

**Figure 5.**
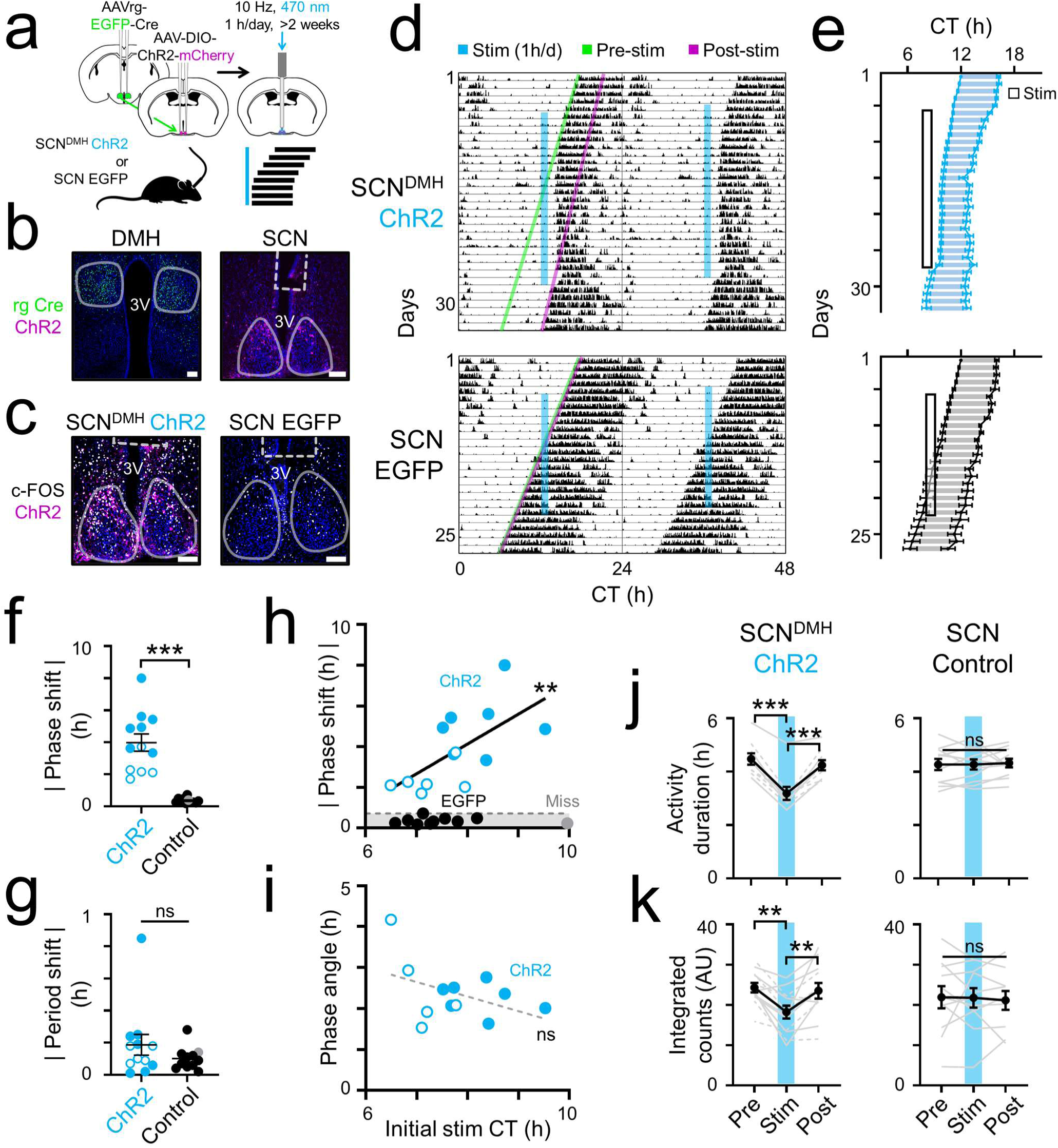
Repeated optogenetic stimulation of SCN^DMH^ neurons entrains locomotor activity rhythms and compresses activity duration. **a)** Schematic of the intersectional optogenetic strategy. AAVrg-EGFP-Cre was injected into the dorsomedial hypothalamus (DMH) and AAV-DIO-ChR2-mCherry was injected into the suprachiasmatic nucleus (SCN) to express channelrhodopsin selectively in DMH-projecting SCN neurons (SCN^DMH^). An optical fiber was implanted above the SCN, and animals received daily optogenetic stimulation (470 nm, 10 Hz, 1 h/day) for more than 2 weeks. **b,c)** Representative histology and stimulation-induced c-FOS labeling. **b)** Intersectional viral expression and fiber placement. *Left*, DMH showing retrograde EGFP labeling (green). *Right*, SCN showing DIO-ChR2-mCherry expression (magenta) in SCN^DMH^ neurons and the fiber optic tract (white dashed outline). **c)** Representative SCN sections showing c-FOS immunoreactivity after optogenetic stimulation in SCN^DMH^-ChR2 animals (*left*) and SCN EGFP controls (*right*). White outlines indicate the SCN or DMH in each image. 3V, third ventricle. Scale bars = 100 µm. **d)** Representative double-plotted actograms from an SCN^DMH^-ChR2 animal (*top*) and an SCN EGFP control animal (*bottom*). Actograms show locomotor activity in constant darkness (DD) during 5 d before stimulation, 16-23 d of daily optogenetic stimulation, and 5-6 d after stimulation. Blue vertical lines indicate the daily stimulation interval. Green lines project pre-stimulation activity onsets forward, and magenta lines project post-stimulation activity onsets backward. The SCN^DMH^-ChR2 animal entrained to the daily stimulation and free-ran from a shifted phase after stimulation ended, whereas the control animal free-ran through the stimulation interval without an obvious phase shift. **e)** Daily activity onsets and acrophases (means ± SEM) for SCN^DMH^-ChR2 animals (*top*, n = 9) and SCN EGFP control animals (*bottom*, n = 6), plotted in relative circadian time with activity onset on day 1 defined as CT 12 for each animal. Shaded regions between onset and acrophase indicate activity duration, defined here as the interval from activity onset to acrophase. The outlined box indicates the daily stimulation interval. SCN^DMH^-ChR2 animals entrained to the stimulation interval and showed compressed activity duration during stimulation, with activity duration increasing again after stimulation ended. Control animals continued to free-run through the stimulation interval. Split/through experimental animals and controls that did not free-run across the stimulation interval are shown in **Supplementary Figure 4**. **f,g)** Absolute phase shift (**f**) and absolute period shift (**g**) following repeated optogenetic stimulation. Phase shift was calculated as the absolute difference between the extrapolated pre-stimulation phase and the back-extrapolated post-stimulation phase, and period shift as the absolute difference between pre- and post-stimulation free-running periods. SCN^DMH^-ChR2 animals (blue, n = 12) were compared with EGFP controls (black, n = 10) and a missed-injection control (gray, n = 1). Filled blue circles indicate bilateral DMH-targeted retrograde labeling; open blue circles indicate unilateral DMH-targeted labeling. Absolute phase shift was significantly greater in SCN^DMH^-ChR2 animals (**f**; Welch’s t-test, p < 0.0001), whereas absolute period shift did not differ between groups (**g**; Welch’s t-test, p = 0.206). **h)** Relationship between initial stimulation time, defined as the circadian time of the first stimulation day, and absolute phase shift. Blue circles indicate SCN^DMH^-ChR2 animals (filled, bilateral DMH hits; open, unilateral DMH hits); black circles indicate EGFP controls; gray circle indicates the missed-injection control. The dashed horizontal line indicates the maximum phase shift observed in any control animal. In ChR2 animals, initial stimulation time was positively correlated with phase shift (simple linear regression, r^2^ = 0.422, p = 0.016). **i)** Relationship between initial stimulation time and phase angle of entrainment, defined as the interval between stimulation onset and the entrained activity rhythm. Filled blue circles indicate bilateral DMH hits, and open blue circles indicate unilateral DMH hits. The mean phase angle of entrainment was 2.37 ± 0.2 h (mean ± SEM) and was not significantly correlated with initial stimulation time (simple linear regression, r^2^ = 0.187, p = 0.160). **j,k)** Activity duration (**j**) and integrated locomotor activity counts (**k**) before, during, and after repeated optogenetic stimulation in SCN^DMH^-ChR2 animals (*left*; **j**, n = 9; **k**, n = 12) and controls (*right*, n = 11, including 10 SCN EGFP controls and 1 missed-injection control). Gray lines connect individual animals; for SCN^DMH^-ChR2 animals, dashed lines indicate unilateral DMH-targeted labeling and solid lines indicate bilateral DMH-targeted labeling. Black circles indicate mean ± SEM. Split/through SCN^DMH^-ChR2 animals were excluded from **j** because activity duration could not be reliably quantified during stimulation, but were included in **k** because integrated counts could be quantified. Activity duration was significantly reduced during stimulation in SCN^DMH^-ChR2 animals, but not controls (two-way repeated-measures ANOVA, significant main effect of time and time × group interaction, p < 0.0001 for both; post hoc Tukey’s multiple comparisons test). Integrated counts were lower during the stimulation epoch than during the pre- or post-stimulation epochs in SCN^DMH^-ChR2 animals, whereas controls showed no significant change across epochs (mixed-effects analysis, significant main effect of time, p = 0.0035, and time × group interaction, p = 0.0433; post hoc Tukey’s multiple comparisons test). **, p < 0.01; ***, p < 0.001; ns, not significant.

In SCN^DMH^-ChR2 animals, repeated daily optogenetic stimulation imposed a new phase relationship between locomotor activity rhythms and the stimulation interval. During stimulation, SCN^DMH^-ChR2 animals progressively entrained to the daily stimulation interval at a stable phase angle. After stimulation ended, they resumed free-running from the stimulation-imposed phase rather than from the phase predicted by their pre-stimulation trajectory. In contrast, SCN EGFP controls continued to free-run through the stimulation interval and remained aligned with their projected pre-stimulation trajectory after stimulation ended (**Fig. 5d,e**). Additional EGFP controls whose free-running trajectories did not cross the stimulation interval during the recording window also showed no obvious post-stimulation phase displacement (**Supplementary Fig. 4a,b**). SCN^DMH^-ChR2 animals had significantly larger absolute phase shifts than controls (Welch’s t-test, p < 0.0001; **Fig. 5f**), whereas absolute period shift did not differ between groups (Welch’s t-test, p = 0.206; **Fig. 5g**). Thus, repeated SCN^DMH^ stimulation displaced the phase of the locomotor activity rhythm without detectably changing the free-running period before versus after stimulation.

We next examined how entrainment dynamics related to each animal’s pre-stimulation rhythm and the circadian time of stimulation. Some SCN^DMH^-ChR2 animals showed a response we term “split/through,” in which one activity component continued to free-run through the stimulation interval while a second component entrained to the daily stimulation (**Supplementary Fig. 4a,c**). Across SCN^DMH^-ChR2 animals, days to entrainment were shorter when stimulation began later in circadian time and longer in animals with longer pre-stimulation free-running periods (for split/through animals, days to entrainment were calculated from the entrained activity component; **Supplementary Fig. 4d,e**). Split/through animals had modestly later initial stimulation times than animals that entrained with a single dominant activity rhythm, although this subgroup was small (**Supplementary Fig. 4f**). Consistent with these timing relationships, initial stimulation time was positively correlated with absolute phase shift, indicating that the size of the post-stimulation phase displacement depended on when daily stimulation first intersected the animal’s ongoing free-running rhythm (simple linear regression, r^2^ = 0.422, p = 0.016; **Fig. 5h**). Once animals entrained, however, the phase angle of entrainment was similar across animals and was not significantly correlated with initial stimulation time (mean phase angle, 2.37 ± 0.2 h; r^2^ = 0.187, p = 0.160; **Fig. 5i**).

Post hoc histology showed that AAVrg-Cre had been injected into both DMH nuclei in some SCN^DMH^ChR2 animals and into one DMH in others, creating an opportunity to ask whether bilateral DMH targeting was necessary for entrainment. Animals with unilateral DMH AAVrg-Cre injection still entrained to stimulation, although animals with bilateral DMH AAVrg-Cre injection showed larger absolute phase shifts. Phase angle of entrainment and absolute period shift did not differ significantly between unilateral and bilateral DMH injection groups (**Supplementary Fig. 4g-i**).

Repeated SCN^DMH^ stimulation also changed the temporal structure of locomotor activity during entrainment. SCN^DMH^-ChR2 animals showed compressed activity duration during stimulation, followed by a return to baseline after stimulation ended (**Fig. 5e**). Because activity offsets are difficult to define reliably in mice, we defined activity duration as the interval from activity onset to acrophase [41]. Activity duration was significantly reduced during stimulation in SCN^DMH^-ChR2 animals but not controls (two-way repeated-measures ANOVA, significant main effect of time and time × group interaction, p < 0.0001 for both; **Fig. 5j**). Because split/through animals did not have a single assignable activity duration during stimulation, we used integrated locomotor activity counts as a complementary measure that could be applied to the full SCN^DMH^-ChR2 cohort. Integrated counts were lower during the stimulation epoch than during the pre- or post-stimulation epochs in SCN^DMH^-ChR2 animals, whereas controls showed no significant changes across epochs (mixed-effects analysis, significant main effect of time, p = 0.034, and time × group interaction, p = 0.043; post hoc Tukey’s multiple comparisons test for SCN^DMH^-ChR2 animals, p < 0.05; **Fig. 5k**). This pattern was similar when SCN^DMH^-ChR2 animals were separated by unilateral or bilateral DMH AAVrg-Cre injection. Both activity duration and integrated counts changed across the stimulation paradigm, but neither differed significantly between injection groups (**Supplementary Fig. 4j,k**). Together, these results show that repeated activation of SCN^DMH^ neurons is sufficient to entrain locomotor activity rhythms, produce persistent phase shifts after stimulation ends, and transiently compress the duration of activity.

## DISCUSSION

This study identifies a direct SCN-to-DMH output population capable of organizing the timing of locomotor activity. We first found that DMH neuronal ablation disrupted locomotor rhythmicity, whereas DMH *Cry1/2* disruption did not, separating neuronal integrity from local molecular timekeeping as requirements for this behavioral output. This dissociation led us to ask how SCN timing reaches the DMH. We identified a sparse population of DMH-projecting SCN neurons concentrated in the dorsal SCN. These neurons were largely distinct from canonical AVP and VIP populations and enriched in a *Prokr2*/*Vipr2*-expressing transcriptional cluster. Activating this population was sufficient to reset the phase and alter the temporal structure of locomotor activity rhythms.

Previous studies that disrupted the DMH using broad excitotoxic or thermal lesions, cell-type-specific ablation, or silencing approaches have reported greatly reduced or disorganized locomotor activity rhythms [5–12]. These findings support an important role for the DMH in circadian behavioral output, but also illustrate why the mechanism underlying this locomotor phenotype has been difficult to resolve. These approaches were not designed to distinguish whether impaired locomotor rhythmicity reflects the loss of DMH neurons themselves, disruption of local DMH molecular timekeeping, involvement of adjacent hypothalamic regions, or some combination of these factors [13]. Our results refine this interpretation by showing that DMH neuronal ablation and local DMH *Cry1/2* disruption have different effects on locomotor activity rhythms. Thus, for the locomotor output measured here, intact DMH neurons were required, whereas local molecular timekeeping in the DMH was not.

Because SCN timing and intact DMH neurons are each required for circadian locomotor rhythmicity, identifying how SCN output reaches the DMH is essential for understanding how timing information is routed through hypothalamic circuits that organize behavioral rhythms. Prior work has emphasized the SPZ as an important intermediate node between the SCN and DMH, and our data do not exclude SPZ-mediated contributions to circadian behavioral output. However, anatomical evidence has also supported direct SCN-to-DMH projections [16–19,23]. In the present study, we identified DMH-projecting neurons within the SCN, showed that they are molecularly distinct from canonical AVP and VIP populations, and demonstrated that their activation can reorganize circadian locomotor timing. These results argue that direct SCN-to-DMH projections should be included as functional components of SCN output to the DMH.

The transcriptional profile of SCN^DMH^ neurons helps define this projection within the SCN network. SCN^DMH^ neurons showed minimal overlap with AVP and no detectable overlap with VIP. Instead, they were most strongly represented in a cluster marked by *Prokr2* and *Vipr2*. This molecular profile is consistent with the anatomical distribution of SCN^DMH^ neurons, because *Prokr2*- and *Vipr2*-expressing neurons are both enriched in the dorsomedial SCN. The *Prokr2*/*Vipr2* enrichment is notable because both genes belong to receptor systems with established roles in SCN network organization. *Prok2*/*Prokr2* signaling contributes to SCN ensemble timing and behavioral output organization, while *Vip*/*Vipr2* signaling supports synchrony and phase organization across the SCN [27,29,42–49]. Together, the dorsomedial distribution and *Prokr2*/*Vipr2* enrichment of SCN^DMH^ neurons suggest that this projection arises from a receptor-defined SCN population positioned within network pathways that organize circadian timing and behavioral output.

The functional effect of activating this population extended beyond acute changes in activity. Animals entrained to the daily stimulation interval, then resumed free-running from the imposed phase after the stimulation protocol ended, producing a large phase displacement without a detectable change in free-running period. This post-stimulation aftereffect distinguishes phase reorganization from transient masking during stimulation.

That distinction makes the effect of SCN^DMH^ stimulation especially notable given how few neurons were recruited. Prior in vivo optogenetic entrainment studies targeted many more SCN neurons than the SCN^DMH^ population studied here. *Drd1a*-Cre-driven ChR2 activation recruited much of the SCN and could entrain locomotor rhythms with phase relationships that resemble photic entrainment [30]. *Vip*-Cre-driven ChR2 activation targeted canonical VIP neurons, which are far more numerous than the SCN^DMH^ neurons studied here, and entrained locomotor rhythms under some stimulation patterns [50]. In both cases, activity-duration compression was not reported as a feature of entrainment. *Cck*-Cre-driven ChR2 activation also targeted a larger SCN population, but produced phase advances without stable entrainment [51]. In contrast, repeated activation of the sparse SCN^DMH^ population produced stable entrainment, persistent post-stimulation phase displacement, and compression of activity duration.

These comparisons suggest that the effect of SCN stimulation depends strongly on which output channel is recruited, not simply on how many SCN neurons are activated. One explanation is that SCN^DMH^ neurons sit near the output end of the SCN timing network. Broader SCN activation may recruit neurons involved in photic entrainment, intra-SCN coupling, or phase computation. Activation of DMH-projecting SCN neurons may instead engage a more specific output channel through which SCN timing reaches DMH circuits that organize rhythmic behavior. In this framework, a small number of SCN^DMH^ neurons can produce a large behavioral effect because they are positioned to route timing information out of the SCN.

A separate feature of the SCN^DMH^ stimulation phenotype was compression of the active phase during entrainment, suggesting that this pathway affects both the phase and temporal structure of activity. In classical circadian models, photoperiod-dependent compression and decompression of daily activity time reflect differential shifts in activity onset and offset, consistent with coupled evening and morning oscillator organization [52,53]. This compression raises the possibility that SCN^DMH^ activation changes how locomotor activity is arranged within the active phase, rather than simply shifting the rhythm earlier or later. Determining whether this reflects differential effects on activity onset and offset, altered DMH control of arousal state, or changes in SCN subnetwork coordination will require experiments that resolve activity timing and circuit dynamics more directly.

The present experiments establish that repeated activation of SCN^DMH^ neurons is sufficient to organize circadian locomotor behavior, but they do not determine whether this population normally carries circadian timing information or is required for endogenous locomotor rhythmicity. Addressing these questions will require defining how SCN^DMH^ neurons vary across circadian time and whether selective loss of this projection disrupts behavioral rhythms. A separate challenge is following this pathway beyond the DMH. This will require identifying which DMH neurons receive SCN^DMH^ input, where those neurons project, and how those downstream pathways shape arousal and locomotor output. Together, these experiments would extend the circuit defined here from direct SCN-to-DMH output to the downstream pathways that structure behavior across circadian time.

## METHODS

### Animals and housing

Male and female wild-type and H11^LSL-Cas9^ mice [34], 2 to 4 months old, on a C57BL/6J background were used for experiments. Mice were singly housed at constant temperature (∼23°C) and humidity (∼40-45%) under a 12 h:12 h light:dark cycle (LD; lights on, 07:00, defined as zeitgeber time 0), with food and water available ad libitum. Mice were transferred to constant darkness (DD) for free-running locomotor activity recordings and optogenetic stimulation experiments as described below. All procedures were approved by the Texas A&M University Institutional Animal Care and Use Committee and performed in accordance with institutional guidelines.

### Stereotaxic surgery

Mice were anesthetized with isoflurane (4% induction at 400 mL/min; 2% maintenance at 30 mL/min) and received buprenorphine-SR (1 mg/kg, s.c.) before surgery. Viral injections were delivered bilaterally unless otherwise specified. All coordinates are reported relative to bregma. Standard injection coordinates were AP - 0.46 mm, ML ±0.16 mm, DV −5.67 mm for the SCN and AP −1.80 mm, ML ±0.30 mm, DV −5.22 mm for the DMH. All viruses were obtained from the University of Zurich Viral Vector Core. Viruses were infused at 1 nL/s, and the injection needle was left in place for 10 min before withdrawal. Mice recovered in their home cages for at least 2 weeks before behavioral experiments.

For neuronal ablation experiments, we used region-targeted viral delivery to ablate neurons in either the SCN or DMH. For SCN ablation, each side received 50 nL of a 1:10 mixture of AAVDJ/8-hSyn-EGFP-Cre (4.80 × 10^12^ GC/mL) and AAVPHP.eB-Ef1a-DO-mCherry-DIO-DTA (7.10 × 10^12^ GC/mL). For DMH ablation, each side received 200 nL of a 1:10 mixture of AAVDJ/8-hSyn-EGFP-Cre (4.80 × 10^12^ GC/mL) and AAVDJ-Ef1a-DIO-Casp3 (4.10 × 10^12^ GC/mL).

For local *Cry1*/*2* disruption experiments, we used region-targeted viral delivery in H11^LSL-Cas9^ mice to disrupt molecular clock function in either the SCN or DMH. Each side received 200 nL of a 1:1 mixture of AAVDJ/8-hSyn-EGFP-Cre and AAVDJ/8-U6-sgCry1/2-mCherry (6.80 × 10^12^ GC/mL) [35]. Control mice received AAVDJ/8-hSyn-EGFP-Cre mixed 1:1 with AAVDJ/8-U6-sgLacZ-mCherry (7.00 × 10^12^ GC/mL).

For retrograde tracing of DMH-projecting SCN neurons, AAVrg-hSyn-EGFP was injected unilaterally into the DMH. Each mouse received 50 nL of AAVrg-hSyn-EGFP (8.50 × 10^12^ GC/mL).

For intersectional tracing of SCN-to-DMH terminals, mice received unilateral injections into the DMH and ipsilateral SCN. The DMH was injected with 50 nL of AAVrg-hSyn-mCherry-iCre (8.40 × 10^12^ GC/mL). The ipsilateral SCN was injected with 75 nL of AAV8-hSyn-DIO-Syp-EGFP (2.50 × 10^12^ GC/mL).

For intersectional optogenetic activation of DMH-projecting SCN neurons, experimental mice received unilateral injections into the DMH and ipsilateral SCN. The DMH was injected with 50 nL of AAVrg-hSyn-EGFP-iCre (6.10 × 10^12^ GC/mL). The ipsilateral SCN was injected with 50 nL of AAVPHP.eB-Ef1a-DIO-ChR2-mCherry (1.40 × 10^13^ GC/mL). Control mice received a unilateral SCN injection of 50 nL AAV9-hSyn-EGFP (2.90 × 10^13^ GC/mL). In both experimental and control mice, a fiber optic cannula (6 mm length, 400 μm core diameter, 0.39 NA; RWD Life Science) was implanted above the SCN at AP −0.45 mm, ML 0 mm, and DV −5.30 mm. Implants were secured with adhesive cement (Parkell) and opaque black dental cement, then covered with a thin layer of black nail polish.

### Wheel-running and locomotor activity recording

Mice were singly housed in cages equipped with running wheels, with food and water available ad libitum. Wheel-running activity was recorded continuously in 6 min bins using ClockLab software (Actimetrics). For neuronal ablation experiments, mice were recorded for at least 2 d in LD followed by at least 12 d in DD. For *Cry1/2* disruption experiments, mice were recorded for at least 7 d in DD before surgery and at least 14 d in DD after surgery. For optogenetic stimulation experiments, mice were recorded for at least 5 d in DD before stimulation, at least 16 d during stimulation, and at least 6 d after stimulation ended. Animals were excluded from analysis if wheel-running data were missing for multiple consecutive days because of equipment failure.

### Optogenetic stimulation

Optogenetic stimulation was performed in DD after at least 5 d of baseline wheel-running recording using established methods [40]. Experimental and control mice were connected to fiber optic patch cables, and light was delivered through the implanted SCN fiber using a high-power 470 nm LED controlled by an LED driver (Thorlabs). Stimulation (470 nm, 10 Hz, 10 ms pulse width) was delivered once per day at 12:00 local clock time for 1 h. Light power was ∼9 mW at the fiber tip. Control mice underwent the same light-delivery protocol.

### Tissue collection and histology

For *Cry1/2* disruption experiments, mice were returned to LD for at least 5 d, or until re-entrained, and tissue was collected at ZT 6. For anatomical tracing experiments, tissue was collected around ZT 6. For neuronal ablation experiments, tissue was collected around 12:00 local clock time. For optogenetic stimulation experiments, mice were entrained to a reverse LD cycle for at least 5 d, or until entrained. These mice then received 30 min of optogenetic stimulation at 12:00 local clock time, and tissue was collected 45 min after stimulation ended.

### Immunohistochemistry

Free-floating sections were processed for immunostaining using established methods [14]. For AVP and VIP overlap experiments, mice received intracerebroventricular colchicine (2 µL; 1 µg/µL) 48 h before perfusion to enhance neuropeptide labeling. Primary antibodies were rabbit anti-CRY2 (1:500; Cry21-A, Alpha Diagnostics), rabbit anti-AVP (1:1000; 20069, Immunostar), guinea pig anti-VIP (1:500; 443 005, Synaptic Systems), rabbit anti-c-FOS (1:2000; 226 008, Synaptic Systems), and chicken anti-mCherry (1:1000; MCHERRY-0020, Aves Labs). Secondary antibodies were CF647 goat anti-rabbit (1:500; 20282, Biotium), CF568 goat anti-guinea pig (1:500; 20492, Biotium), and CF568 goat anti-chicken (1:500; 20104, Biotium). Sections were counterstained with DAPI where appropriate. All brains within each staining experiment were processed in parallel using identical antibody concentrations, incubation durations, and wash conditions. Sections were imaged on a Nikon Ti2-E equipped with an X-Light V3 spinning-disk confocal (Crest Optics), using acquisition settings held constant across animals within each analysis.

### Histological quantification and anatomical mapping

For ablation and CRY2 validation, target regions were identified using DAPI staining patterns and atlas landmarks. For neuronal ablation experiments, DAPI optical density within the injected SCN or DMH was measured in ImageJ and compared with the surrounding hypothalamus. Ablation cases were classified as target hits when DAPI optical density within the injected region was reduced by ≥75% relative to the surrounding hypothalamus; cases not meeting this criterion were classified as misses. For *Cry1/2* disruption experiments, SCN and DMH ROIs were defined from DAPI staining. Within each ROI, CRY2-positive cells were identified in ImageJ using identical threshold settings across sections followed by particle analysis. CRY2-positive cell counts were quantified across rostral-to-caudal sections for each animal and compared between sgCry1/2- and sgLacZ-injected mice.

For anatomical mapping of DMH-projecting SCN neurons, EGFP-positive cells were manually counted in tissue from retrograde tracing experiments. Cells were counted bilaterally across 10 to 12 sections spanning the rostral-caudal extent of the SCN for each animal. SCN boundaries were defined by the sharp transition from the densely packed DAPI-labeled nuclei of the SCN to the lower-density surrounding hypothalamus, together with landmarks from the Paxinos and Franklin mouse brain atlas and the Allen Mouse Brain Reference Atlas. For SCN-versus-SPZ comparisons, the SPZ was operationally defined as the tissue immediately dorsal to the SCN on each section in which the SCN was present. This definition was used to provide an inclusive estimate of retrogradely labeled cells in the SCN-adjacent SPZ region. EGFP-positive cells outside the SCN and the operationally defined SPZ were noted separately but were not included in SCN-versus-SPZ counts. Counts were summarized per animal as the total number of EGFP-positive cells assigned to the SCN or SPZ.

For AVP and VIP overlap analysis, EGFP-positive DMH-projecting SCN neurons were scored for colocalization with AVP or VIP immunolabeling in colchicine-treated tissue. For each 40 µm SCN section, z-stacks were analyzed plane-by-plane to assess overlap within individual cell bodies. Labeled cells were manually counted across individual z-planes, and EGFP-positive cells were classified as overlapping with AVP or VIP only when the same cell body could be followed across adjacent optical sections. This approach was used to distinguish true overlap from close apposition of neighboring labeled cells. Counts were summarized per animal across the rostral-caudal extent of the SCN, with AVP and VIP results reported separately.

For intersectional tracing and optogenetic verification, tissue sections were examined histologically for viral expression, injection placement, terminal labeling, and fiber placement. In terminal tracing experiments, DMH injection placement, mCherry-iCre expression in the SCN, and Syp-EGFP-labeled terminals in the DMH were evaluated from the same tissue series. In optogenetic experiments, ChR2-mCherry expression in the SCN and fiber tract placement were evaluated histologically. c-FOS immunostaining after light delivery was used as a qualitative measure of optogenetic activation in the stimulated SCN region.

### Single-cell RNA sequencing

For scRNA-seq tissue collection and cell isolation, 10 mice received bilateral DMH injections of AAVrg-EGFP, and SCN tissue was collected around ZT 6. Mice were euthanized and brains were rapidly dissected into carbogenated ACSF. For each animal, both SCNs were manually microdissected from 1-2 coronal sections cut at 300 µm on a vibratome (Campden Instruments). Dissected SCN tissue from all animals was pooled and dissociated using papain-based enzymatic digestion followed by mechanical trituration. Dissociated cells were fixed overnight at 4°C using the 10x Genomics Flex Sample Preparation v2 Kit, and DAPI-positive fixed cells were isolated by FACS.

Fixed cells were prepared for scRNA-seq using the 10x Genomics GEM-X Flex Gene Expression Mouse Kit. Cells were divided into four portions and hybridized with whole-transcriptome probe sets containing unique sample multiplexing barcodes and custom EGFP probes. After hybridization, samples were pooled and processed for GEM generation and library preparation by the Texas A&M Molecular Genomics Core. Libraries were sequenced on an Illumina NextSeq 2000 using a P3 flow cell with approximately 1.2 × 10^9^ reads.

Reads were aligned to the GRCm39 mouse reference genome using Cell Ranger through the 10x Genomics Cloud analysis interface. Custom EGFP probes were included to enable detection of retrograde tracer-derived EGFP transcripts. Filtered count matrices were imported into RStudio and analyzed using Seurat v5. Cells with >10% mitochondrial gene content were excluded, and cells with 1,000-8,000 detected genes were retained for downstream analysis. Putative SCN clusters were curated using canonical SCN marker expression, including *Avp*, *Vip*, *Cck*, *Nms*, *Vipr2*, *Grp*, and *Prokr2*, together with removal of non-SCN hypothalamic clusters marked by *Sst* or *Trh*.

The remaining putative SCN population was re-normalized, scaled, and re-embedded using PCA, clustering, and UMAP. EGFP-positive cells were identified by summing normalized expression across the EGFP-1, EGFP-2, and EGFP-3 transgene entries, with cells classified as EGFP-positive when summed EGFP expression was greater than zero. Putative SCN clusters were grouped into transcriptionally related groups using hierarchical clustering of z-scored mean expression profiles across variable genes, yielding nine transcriptional groups. Group marker genes were identified with Seurat’s FindAllMarkers function, and EGFP-positive cell distribution across groups was calculated as the percentage of all EGFP-positive neurons assigned to each group.

### Locomotor rhythm analysis

Wheel-running activity was analyzed in ClockLab using established methods [30]. Free-running period was estimated from DD records using Lomb-Scargle periodogram analysis. For neuronal ablation and *Cry1/2* disruption experiments, locomotor rhythmicity was quantified using Cosinor amplitude.

Activity onsets were calculated in ClockLab and used to estimate rhythm phase. For optogenetic stimulation experiments, entrainment was defined as the first day on which activity onsets adopted a stable phase relationship to the daily stimulation interval, with <15 min of day-to-day variation. Phase shifts were calculated in ClockLab by comparing the extrapolated pre-stimulation onset regression line with the post-stimulation onset regression line.

To quantify changes in the temporal distribution of locomotor activity, we measured the interval from daily activity onset to acrophase, calculated in ClockLab. For animals that developed two distinct daily activity components during stimulation, standard ClockLab onset and acrophase values were used until the components separated. After separation, the earlier component was defined by the ClockLab-detected activity onset and the center of mass of the pre-stimulation bout. The later, stimulation-entrained component was defined by manually identified activity onset and the center of mass of the post-stimulation bout.

### Statistical analysis

Statistical analyses were performed in GraphPad Prism. Data are presented as mean ± SEM unless otherwise indicated. Individual animals were used as the unit of biological replication unless otherwise stated. Statistical significance was defined as p < 0.05. For comparisons between two groups, paired or unpaired t-tests were used as appropriate for the experimental design. Welch’s correction was used when equal variances were not assumed. One-sample t-tests were used to compare normalized values against a defined reference value. One-way or two-way ANOVA was used for multi-group or multi-factor comparisons, with repeated-measures designs used when measurements were repeated within the same animals. Mixed-effects analysis was used for repeated-measures datasets with missing values. Fisher’s LSD, Tukey’s, or Sidak’s multiple-comparisons tests were used for post hoc comparisons as indicated in the figure legends. Relationships between stimulation timing and behavioral outcomes were evaluated using simple linear regression.

## DATA AVAILABILITY

All data supporting the findings of this study are available within the article and its Supplementary Material.

## ACKNOWLEDGMENTS

This work was supported by National Institutes of Health Grant R35GM151020 (J.R.J.) and a research grant from the Whitehall Foundation (J.R.J.). We thank members of the Jones Lab for helpful discussions and feedback on the manuscript.

## Notes

### Competing Interest Statement

The authors have declared no competing interest.

