## Supplementary Figures for "A direct SCN-to-DMH output pathway organizes circadian behavioral timing"

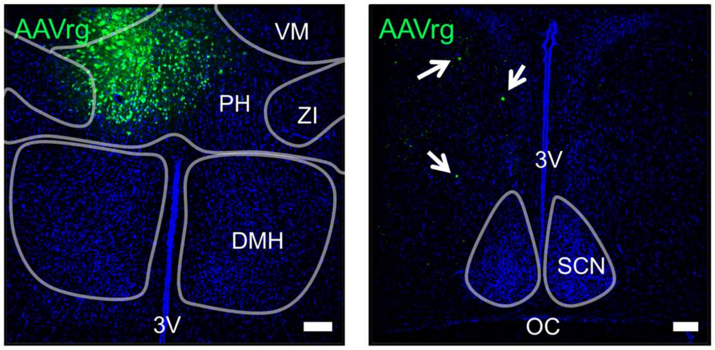


**Supplementary Figure 1. Dorsal off-target DMH injections do not label SCN neurons.** *Left*, representative image of an AAVrg-Syn-EGFP injection centered dorsal to the DMH. White outlines indicate the DMH, posterior hypothalamus (PH), zona incerta (ZI), and ventromedial thalamic nucleus (VM). EGFP labeling is concentrated primarily in the PH, with limited spread into the ZI and VM. *Right*, representative image of retrograde EGFP labeling in the SCN region following a dorsal off-target injection. No EGFP-positive cells were observed in the SCN. White outline indicates the SCN. White arrows indicate labeled cells dorsal to the SCN, likely in the subparaventricular zone. 3V, third ventricle; OC, optic chiasm. Scale bars = 100 µm.

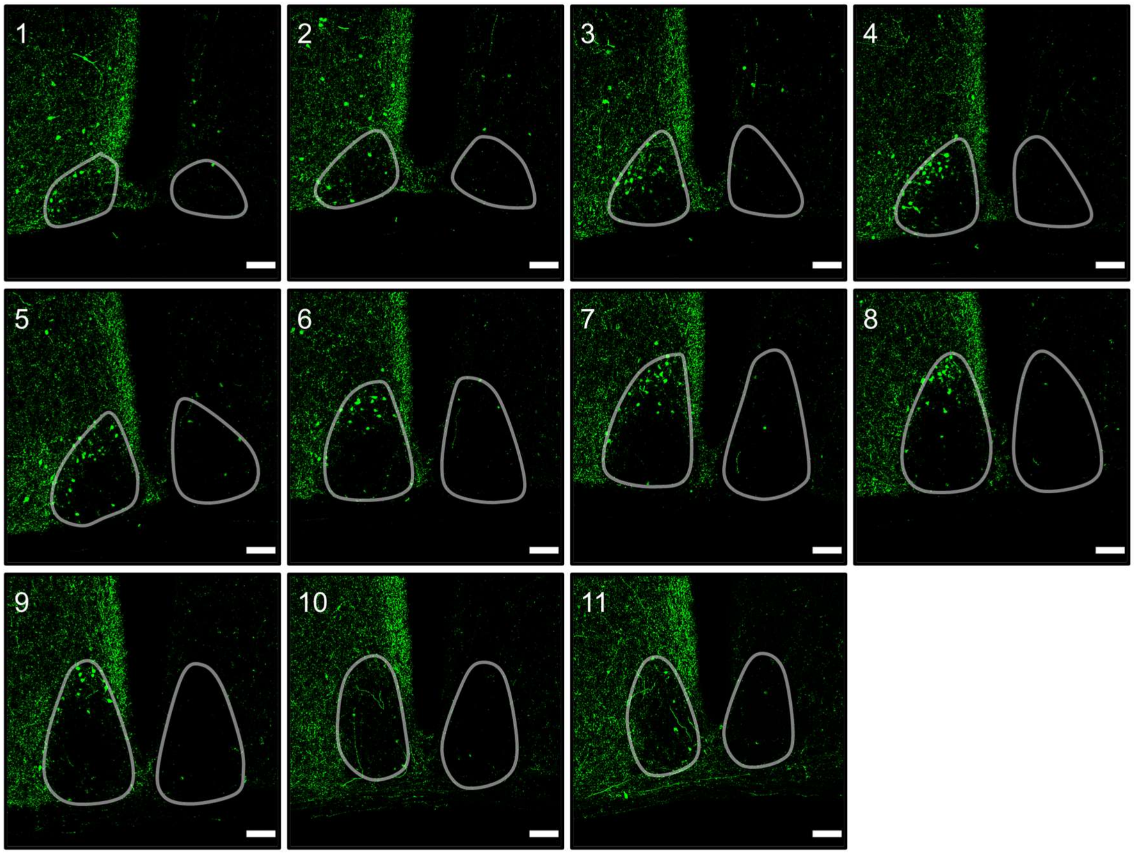


**Supplementary Figure 2. Rostral-to-caudal distribution of retrogradely labeled SCN$^{DMH}$ neurons across the SCN.** Representative 40 μm coronal sections spanning the rostral-to-caudal extent of the SCN (sections 1-11) after unilateral AAVrg-Syn-EGFP injection into the DMH. EGFP-positive SCN$^{DMH}$ neurons are shown in green. White outlines indicate the SCN, defined by DAPI cell density relative to surrounding tissue. Most labeled cells were located in the SCN across the series. A small number of labeled cells dorsal and lateral to the SCN were observed in the most rostral sections, likely corresponding to medial preoptic area and/or median preoptic nucleus neurons that also project to the DMH. Scale bars = 100 μm.

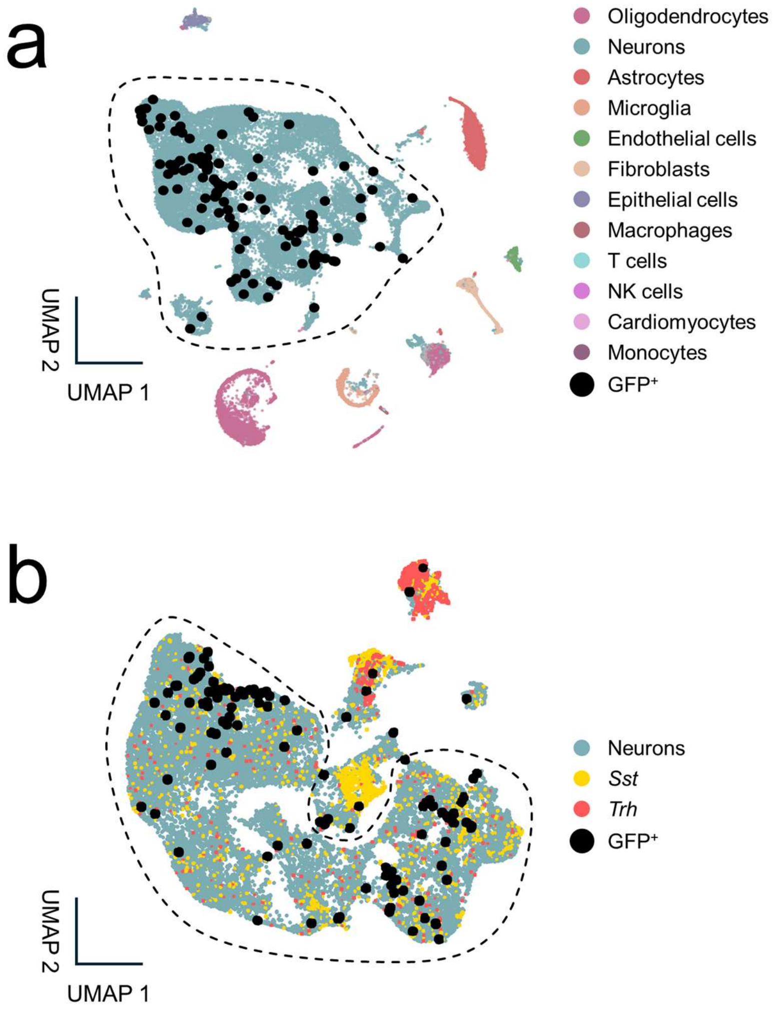


**Supplementary Figure 3. Manual curation of putative SCN neurons for scRNA-seq analysis.**

**a)** UMAP of all sequenced cells colored by major cell class identity. EGFP-positive retrogradely labeled DMH-projecting cells are shown as black dots and localized within the neuronal cluster. Dashed outline indicates the neuronal cluster selected for further analysis. **b)** UMAP of the isolated neuronal cluster showing refinement of putative SCN neurons. EGFP-positive retrogradely labeled DMH-projecting neurons are shown as black dots. Neurons enriched for somatostatin (*Sst*, yellow) or thyrotropin-releasing hormone (*Trh*, red), markers of non-SCN hypothalamic populations, formed distinct UMAP islands outside the main cluster and were excluded from further analysis. Dashed outline indicates the remaining population selected as putative SCN neurons.

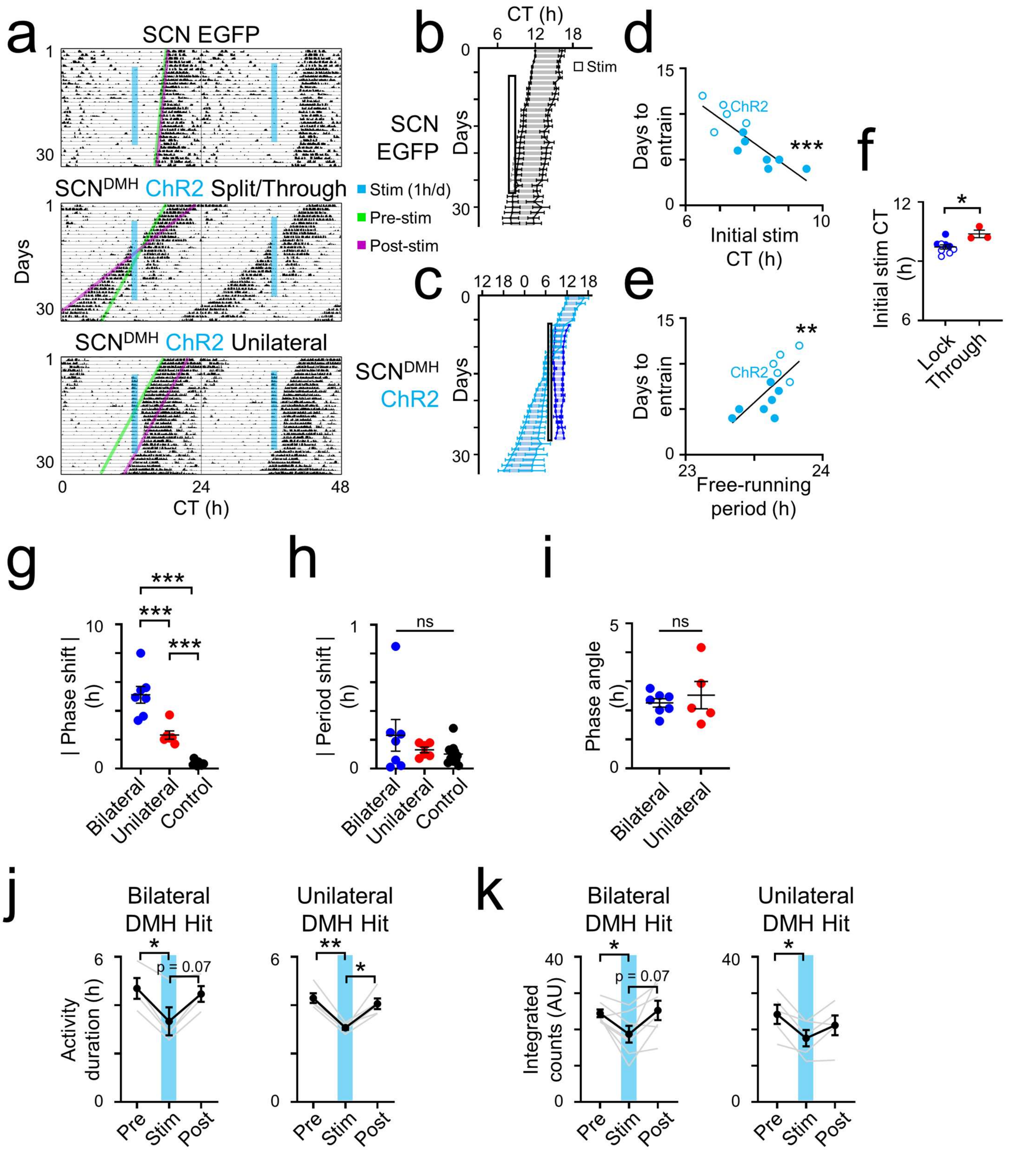


**Supplementary Figure 4. Additional entrainment patterns and unilateral/bilateral response comparisons during repeated optogenetic stimulation of SCN$^{DMH}$ neurons. a)** Representative double-plotted actograms illustrating additional control, split/through, and unilateral response patterns. *Top*, an SCN EGFP control animal with a longer free-running period, showing continued free-running without an obvious phase shift. Because its activity rhythm did not cross the stimulation interval during the recording window, this example is less informative for assessing phase resetting by stimulation. *Middle*, a "split/through" $SCN^{DMH}$-ChR2 animal in which the primary activity rhythm split into two components, one that continued to free-run and a second that locked to the stimulation interval. *Bottom*, an $SCN^{DMH}$-ChR2 animal with unilateral DMH-targeted retrograde labeling, showing entrainment to the daily stimulation interval and a phase-shifted free run after stimulation ended. Actograms show locomotor activity in constant darkness before, during, and after optogenetic stimulation. Blue vertical lines indicate the daily stimulation interval. Green lines project pre-stimulation activity onsets forward, and magenta lines project post-stimulation activity onsets backward. **b,c)** Daily activity onsets and acrophases (mean ± SEM) for SCN EGFP control animals whose free-running trajectories did not cross the stimulation interval during the recording window (**b**, n = 5) and split/through $SCN^{DMH}$-ChR2 animals (**c**, n = 3). In **c**, the primary activity component (light blue) corresponds to daily activity onset and acrophase or, on days with two distinct bouts, the center of mass of the pre-stimulation bout; the secondary entrained component (dark blue) corresponds to activity onset and center of mass of the post-stimulation bout. Shaded regions indicate activity duration as defined in **Figure 5**, and the outlined box indicates the daily stimulation interval. **d,e)** Relationship of days to entrainment with initial stimulation time (**d**) or pre-stimulation free-running period (**e**) in $SCN^{DMH}$-ChR2 animals (n = 12; filled circles, bilateral DMH hits; open circles, unilateral DMH hits). For split/through animals, days to entrainment were calculated from the secondary entrained activity component. Days to entrainment were defined as the number of days required to reach a stable phase angle relative to the daily stimulation interval. Initial stimulation time was negatively correlated with days to entrainment (**d**; simple linear regression, $r^2 = 0.729$, $p = 0.001$), whereas pre-stimulation free-running period was positively correlated with days to entrainment (**e**; simple linear regression, $r^2 = 0.501$, $p = 0.010$). f) Initial stimulation time in $SCN^{DMH}$-ChR2 animals that locked to the stimulation interval ("lock," n = 9; open circles, unilateral DMH-targeted labeling, n = 5; filled circles, bilateral DMH-targeted labeling, n = 4) and animals that exhibited split/through responses ("through," n = 3). Initial stimulation time was modestly later in split/through animals than in lock animals (lock, 7.45 ± 0.22 h; split/through, 8.77 ± 0.38 h; mean ± SEM; Welch's t-test, p = 0.048). **g,h)** Absolute phase shift (**g**) and absolute period shift (**h**) following repeated optogenetic stimulation in $SCN^{DMH}$-ChR2 animals with bilateral DMH-targeted labeling (dark blue, n = 7), unilateral DMH-targeted labeling (red, n = 5), and controls (black, n = 11, including 10 SCN EGFP controls and 1 missed-injection control). Absolute phase shift differed significantly across groups (**g**; one-way ANOVA with post hoc Tukey's multiple comparisons test, $p < 0.0001$), whereas absolute period shift did not (**h**; one-way ANOVA with post hoc Tukey's multiple comparisons test, $p > 0.05$). **i)** Phase angle of entrainment in $SCN^{DMH}$-ChR2 animals with bilateral DMH-targeted labeling (dark blue, n = 7) or unilateral DMH-targeted labeling (red, n = 5). Phase angle of entrainment did not differ significantly between bilateral and unilateral animals (Welch's t-test, p = 0.6110). **j,k)** Activity duration (**j**) and integrated locomotor activity counts (**k**) before, during, and after repeated optogenetic stimulation in $SCN^{DMH}$-ChR2 animals with bilateral DMH-targeted labeling (*left*; **j**, n = 4; **k**, n = 7) or unilateral DMH-targeted labeling (*right*; n = 5 for both panels). Gray lines connect individual animals, and black circles indicate mean ± SEM. Split/through animals were excluded from **j** because activity duration could not be reliably quantified during stimulation, but were included in **k** because integrated counts could be quantified. Both measures were significantly altered across the stimulation paradigm (**j**, two-way repeated-measures ANOVA with post hoc Tukey's multiple comparisons test, main effect of time, $p < 0.0001$; **k**, two-way repeated-measures ANOVA with Sidak's multiple comparisons test, main effect of time, $p = 0.0009$), and did not differ significantly between bilateral and unilateral groups. *, $p < 0.05$; **, $p < 0.01$; ***, $p < 0.001$; ns, not significant.
